# Ventriculo-arterial uncoupling is associated with VO_2_ dependency in cardiac surgical patients

**DOI:** 10.1101/602607

**Authors:** Pierre-Grégoire Guinot, Maxime Nguyen, Pierre Huette, Osama Abou-Arab, Belaid Bouhemad, Dan Longrois

**Author notes:** Correspondence to: Pr Pierre-Grégoire Guinot, Anaesthesiology and Critical Care Department, Dijon University Hospital, 2 Bd Maréchal de Lattre de Tassigny, F-21000 Dijon, France., Twitter: @guinot pg.

## Abstract

**Background:** The clinical relevance of V-A (un)coupling in critically ill patients is under investigation. In this study we measured the association between V-A coupling and oxygen consumption (VO_2_) response in patients with acute circulatory instability following cardiac surgery.

**Methods and results:** Sixty-one cardio-thoracic ICU patients who received fluid challenge or norepinephrine infusion were included. Arterial pressure, cardiac output (CO), heart rate (HR), arterial (E_A_), and ventricular elastances (E_V_), total indexed peripheral resistance (TPRi) were assessed before and after hemodynamic interventions. VO_2_ responders were defined as VO_2_ increase > 15 %. V-A coupling was evaluated by the ratio E_A_/E_V._ Left ventricle stroke work (SW) to pressure volume area (PVA) ratio was calculated. In the overall population, 24 patients (39%) were VO_2_ responders and 48 patients were uncoupled (i.e., E_A_/E_V_ ratio > 1.3): 1.9 (1.6-2.4). Most of the uncoupled patients were classified as VO_2_ responders (28 of 31 patients, p=0.031). Changes in VO_2_ were correlated with those of TPRi, E_A_, E_A_/E_V_ and CO. E_A_/E_V_ ratio predicted VO_2_ increase with an AUC of 0.76 [95 % CI: 0.62-0.87]; p=0.001. In multivariate and principal component analyses, E_A_/E_V_ and SW/PVA ratios were independently associated (P < 0.05) with VO_2_ response following interventions.

**Conclusions:** VO_2_ responders were characterized by baseline V-A uncoupling due to high E_A_ and low E_V_. Baseline E_A_/E_V_ and SW/PVA ratios were associated with VO_2_ changes independently of the hemodynamic intervention used. These results further underline the pathophysiological significance of V-A uncoupling in patients with hemodynamic instability.

## Introduction

Acute circulatory failure following cardiac surgery is characterized by an imbalance between oxygen delivery (DO_2_) and oxygen consumption (VO_2_) which results in tissue hypoxia and organ dysfunction (1). In clinical practice, the difficulty is to identify parameters that are clinically relevant to become endpoints for titration of interventions. Increasing DO_2_ is an accepted goal for optimization following cardiac surgery (2, 3) especially if decreased DO_2_. Thus, predicting VO_2_ responsiveness could identify the patients for which DO_2_ increase is the most beneficial.

The ventriculo-arterial coupling (V-A coupling), describes the interactions between the ventricles and the large arteries from an integrated pressure-volume relationship (4–6). The left ventricle and the arterial system are described by their elastances (ventricular elastance (E_V_), arterial elastance (E_A_)), and V-A coupling is defined by the ratio of E_A_/E_V_ (4). The efficacy and efficiency of the cardiovascular system are the result of regulated interactions between the heart and the vascular system. The optimal hemodynamic intervention in patients with acute circulatory failure would improve efficacy with the lowest energetic cost (high efficiency) for the cardiovascular system (7).

Cardiology studies have demonstrated that V-A coupling may represent a parameter that describes the energetic cost in particular when the left ventricular function is altered (8, 9). There is wide evidence that V-A (un)coupling is a hemodynamic parameter that is associated with patient outcomes (8, 10–13). The relevance of V-A (un)coupling as a parameter of hemodynamic optimization in patients with acute circulatory failure could be related to the fact that V-A (un)coupling is a parameter of cardiovascular efficiency whereas the classical hemodynamic parameters are exclusively parameters of cardiovascular efficacy (2, 3).

In our continuous attempt to investigate the clinical relevance of V-A (un)coupling in critically ill patients we designed the present study in order to analyse the effects of two types of interventions: fluid challenge (FC) or norepinephrine infusion on systemic oxygenation parameters (as indicators of cardiovascular efficacy) and on V-A coupling (as an indicator of cardiovascular efficiency). The main objective of this study was to investigate the relationship between E_A_/E_V_ ratio and changes in VO_2_ upon treatment of hemodynamic instability following cardiac surgery. The second objectives were to compare V-A coupling and oxygenation derivated parameters (central venous saturation (ScVO_2_), gap CO_2_) as predictor of VO_2_ changes following hemodynamic treatment.

## METHODS

### Ethics

Ethical approval for this study (RNI2014-39) was provided by *Comité de Protection des Personnes Nord-Ouest II*, Amiens, France (Chairperson T Bourgueil) on 9 February 2015. All patients received written information and gave their verbal consent to participation. The present manuscript was drafted in compliance with the STROBE checklist for cohort studies (12).

### Patients

This prospective, observational study was performed in the cardiothoracic ICU at University Hospital between 2015 and 2017. The main inclusion criteria were as follows: age 18 or over, controlled positive ventilation, and a clinical decision to treat hemodynamic instability by FC and/or norepinephrine. The indications for FC were arterial hypotension: a systolic arterial pressure (SAP) below 90 mmHg and/or a mean arterial pressure (MAP) below 65 mmHg, and/or stroke volume (SV) variation of more than 10%, and/or clinical signs of hypoperfusion (skin mottling, and a capillary refill time of more than 3 sec). In the present study, FC always consisted of a 10-minute infusion of 500 ml of lactated Ringer’s solution. The indications for norepinephrine were persistent arterial hypotension (SAP less than 100 mmHg and/or MAP less than 65 mmHg) despite FC (13). The non-inclusion criteria were permanent arrhythmia, heart conduction block, the presence of an active pacemaker, poor echogenicity, aortic regurgitation, and right heart failure.

### Measurement and calculations of left ventricular elastance, arterial elastance, and ventriculo-arterial coupling

Stroke volume (SV; mL) and cardiac output (CO; l min^−1^) were calculated by using transthoracic echocardiography (CX50 ultrasound system and an S5-1 Sector Array Transducer, Philips Medical System, Suresnes, France). Mean echocardiographic parameters were retrospectively calculated from five measurements (regardless of the respiratory cycle). E_V_ was estimated at the bedside using the non-invasive single beat method described by Chen *et al*. and validated by conference expert (14, 15). E_A_ was estimated by using the formula E_A_= end-systolic pressure (ESP=0.9 * SAP)/SV (16). SAP was measured by using invasive radial artery catheter. The total energy generated by each cardiac contraction is called the “pressure-volume area” (PVA), which is the sum of the external mechanical work exerted during systole (SW) and the potential energy (PE) stored at the end of systole: PVA = SW + PE (17). The PVA has been demonstrated to be linearly related to myocardial oxygen consumption (17, 18). SW is calculated as ESP × SV. PE is calculated as ESP × ((ESV-V_0_)/2), and assumes that V_0_ is negligible when compared with ESV. We calculated total indexed peripheral resistance (TPRi) as TPRi = MAP-central venous pressure (CVP)/cardiac index (mmHg ml^−1^ m^−2^).

### Oxygenation parameters

We recorded the ventilator settings (tidal volume, plateau pressure and end-expiratory pressure) at baseline. All parameters were measured on arterial and central venous blood gases (supplementary File 1).

### Study procedures

Maintenance or withdraw of preoperative medications followed guidelines. Anaesthesia and cardiopulmonary bypass procedures were standardised for all patients. During the study period, the patients were mechanically ventilated in volume-controlled mode, with a tidal volume set to 7-9 ml kg^−1^ ideal body weight, and a positive end-expiratory pressure (PEEP) of 5-8 cmH_2_O, and sedated with Propofol. Ventilator settings (oxygen inspired fraction, tidal volume, respiratory rate and end positive pressure) were not modified during the study period. The following clinical parameters were recorded: age, gender, weight, ventilation parameters, and primary diagnosis. After an equilibration period, HR, SAP, MAP, diastolic arterial pressure (DAP), CVP, SV, CO, and arterial/venous oxygen content were measured at baseline.

### Statistical analysis

In the absence of preliminary data, we designed an observational study with a convenience sample of 61 consecutive patients. Such size could enable to demonstrate a correlation (0.3 to 0.5) between E_A_/E_v_ ration and VO_2_ response with a power of 0.8 and alpha error of 0.05. The variables’ distribution was assessed using a D’Agostino-Pearson test. Data are expressed as the number, proportion (in percent), mean ± standard deviation (SD) or the median [interquartile range (IQR)], as appropriate. Patients were classified as VO_2_ responders or non-responders as a function of the effect of hemodynamic interventions (FC or norepinephrine) on VO_2_. VO_2_ response was defined as an increase of more than 15% in the VO_2_ (19). The non-parametric Wilcoxon rank sum test, Student’s paired t test, Student’s t test, and the Mann-Whitney test were used to assess statistical significance, as appropriate. Because, we have analysed several correlated hemodynamic and perfusion variables, we performed three different analysis to evaluate the association between ventriculo-arterial coupling and VO_2_: a linear correlation analysis, principal component analysis and predictability analysis. Linear correlations were tested using Pearson’s or Spearman’s rank method. The principal component analysis transforms correlated variables into uncorrelated variables that may explain VO_2_ changes. A principal component analysis was carried out by including fourteen baseline variables. The VO_2_ changes following therapeutics was included as a supplementary variables. A receiver-operating characteristic curve was established for the ability of ScVO_2_ and E_V_/E_A_ ratio to predict an increase of more than 15% in VO_2_. The threshold for statistical significance was set to *p*<0.05. R software (version 3.5.0) with FactoMineR package was used for all statistical analyses.

## RESULTS

Of the 65 included patients, four were excluded (Supplementary file 2), and so the final analysis concerned 61 subjects (Table 1). At baseline 48 patients (78%) were uncoupled with a median E_A_/E_V_ ratio of 1.9 (1.6-2.4) in relation to abnormally low E_V_ (1.1 (0.9-1.6)), as compared to preserved E_A_ (2 (1.5-2.7)). In the overall population, 31 patients (48 %) were classified as VO_2_ responders. The percentage of VO_2_ responders did not differ between the two groups (16 (48%) out of 33 vs 15 (54%) out of 28, p=0.799).

**Table 1.**
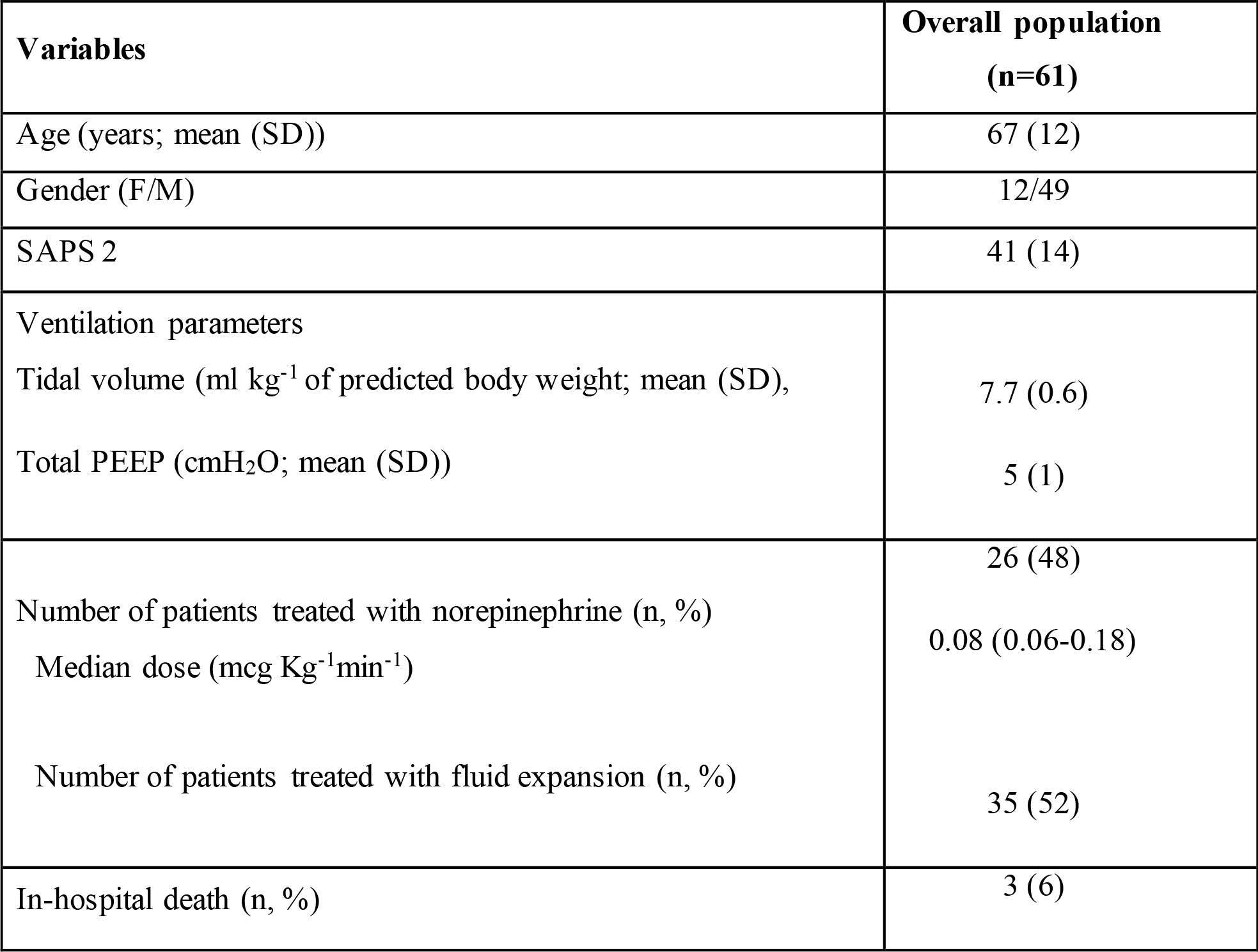
Characteristics of the study participants on inclusion. Values are expressed as the mean or the number (%).

| **Variables** | **Overall population**<br>**(n=61)** |
| --- | --- |
| Age (years; mean (SD)) | 67 (12) |
| Gender (F/M) | 12/49 |
| SAPS 2 | 41 (14) |
| Ventilation parameters |  |
| Tidal volume (ml kg<sup>-1</sup> of predicted body weight; mean (SD), | 7.7 (0.6) |
| Total PEEP (cmH<sub>2</sub>O; mean (SD)) | 5 (1) |
| Number of patients treated with norepinephrine (n, %) | 26 (48) |
| Median dose (mcg Kg<sup>-1</sup>min<sup>-1</sup>) | 0.08 (0.06-0.18) |
| Number of patients treated with fluid expansion (n, %) | 35 (52) |
| In-hospital death (n, %) | 3 (6) |

### Combined analysis of the effects of the two therapeutic interventions on systemic parameters (Table 2, Figure 1)

At baseline, VO_2_ responders had higher E_A_/E_V_ ratio, and lower SW/PVA ratio and VO_2_ than VO_2_ non-responders. Therapeutic interventions increased SAP, MAP, CO and DO_2_ in the overall population. VO_2_ responders were characterized by an increased SW/PVA ratio, and a decreased HR, and TPRi. VO_2_ non-responders were characterized by an increased E_A_, E_V_, ScVO_2_, TPRi, and a decreased gapCO_2_.

**Table 2.** Comparison of haemodynamic parameters in VO_2_ responders and VO_2_ non-responders. Values are expressed as the mean (SD) or the median [interquartile range]. **CO**, cardiac output; ***DAP***, diastolic arterial pressure; ***DO_2_***, oxygen delivery; **FC**, fluid challenge; **HR**, heart rate; **LVEF**, lef ventricular ejection fraction; **MAP**, mean arterial pressure; **SAP**, systolic arterial pressure; **SV**, stroke volume; **TPRi**, total indexed peripheral resistance; ***VO_2_***, oxygen consumption; ^**$**^: *p*<0.05 within groups (pre-/post-FC).

| Hemodynamic variables | VO<sub>2</sub> Responders<br>(n=31) | VO<sub>2</sub> Non-responders<br>(n=30) | *P* value |
| --- | --- | --- | --- |
| **HR (bpm)** |  |  |  |
| Pre | 78 (21) | 82 (19) | 0.410 |
| Post | 75 (20)<sup>\\$</sup> | 80 (16) | 0.274 |
| **SAP (mmHg)** |  |  |  |
| Pre | 99 (19) | 91 (14) | 0.093 |
| Post | 124 (19)<sup>\\$</sup> | 116 (20)<sup>\\$</sup> | 0.333 |
| **MAP (mmHg)** |  |  |  |
| Pre | 68 (15) | 65 (10) | 0.260 |
| Post | 84 (14)<sup>\\$</sup> | 80 (12)<sup>\\$</sup> | 0.707 |
| **DAP (mmHg)** |  |  |  |
| Pre | 52 (13) | 52 (10) | 0.971 |
| Post | 62 (14) | 62 (12) | 0.961 |
| **SV (ml)** |  |  |  |
| Pre | 41 (14) | 46 (18) | 0.413 |
| Post | 57 (19)<sup>\\$</sup> | 50 (16)<sup>\\$</sup> | 0.118 |
| **CO (L min<sup>-1</sup>)** |  |  |  |
| Pre | 3.2 (1.1) | 3.6 (1) | 0.159 |
| Post | 4.1 (1)<sup>\\$</sup> | 3.8 (0.9)<sup>\\$</sup> | 0.357 |
| **TPRi (mmHg ml<sup>-1</sup> m<sup>-2</sup>)** |  |  |  |
| Pre | 42 (14) | 34 (15) | 0.056 |
| Post | 39 (17) <sup>\$</sup> | 39 (14) <sup>\$</sup> | 0.945 |
| **E<sub>A</sub> (mmHg ml<sup>-1</sup>)** |  |  |  |
| Pre | 2.3 (1) | 2.1 (1) | 0.358 |
| Post | 2.2 (0.8) | 2.4 (1.1) <sup>\$</sup> | 0.343 |
| **E<sub>V</sub> (mmHg ml<sup>-1</sup>)** |  |  |  |
| Pre | 1.2 (0.6) | 1.3 (0.6) | 0.252 |
| Post | 1.2 (0.6) | 1.6 (0.7) <sup>\$</sup> | 0.036 |
| **E<sub>A</sub>/E<sub>V</sub>** |  |  |  |
| Pre | 2.2 (0.6) | 1.6 (0.6) | 0.002 |
| Post | 2 (0.9) | 1.6 (0.5) | 0.023 |
| **SW/PVA ratio** |  |  |  |
| Pre | 0.55 (0.12) | 0.62 (0.11) | 0.008 |
| Post | 0.62 (0.15) <sup>\$</sup> | 0.62 (0.12) | 0.891 |
| **DO<sub>2</sub> (ml min<sup>-1</sup>)** |  |  |  |
| Pre | 482 (179) | 504 (146) | 0.603 |
| Post | 635 (219) <sup>\$</sup> | 539 (149) <sup>\$</sup> | 0.047 |
| **VO<sub>2</sub> (ml min<sup>-1</sup>)** |  |  |  |
| Pre | 132 (54) | 180 (53) | 0.001 |
| Post | 198 (61) <sup>\$</sup> | 167 (53) <sup>\$</sup> | 0.041 |
| **ScVO<sub>2</sub> (%)** |  |  |  |
| Pre | 67 (12) | 60 (9) | 0.01 |
| Post | 63 (9) <sup>\$</sup> | 65 (8) <sup>\$</sup> | 0.842 |
| **GapCO<sub>2</sub> (mmHg)** |  |  |  |
| Pre | 9 (4) | 9 (2) | 0.842 |
| Post | 9 (4) | 7 (5) <sup>\$</sup> | 0.061 |
| **Arterial lactate (mmol l<sup>-1</sup>)** |  |  |  |
| Pre | 1.5 (1.3-2.1) | 1.6 (1.3-2.1) | 0.882 |
| Post | 1.5 (1.2-2.1) | 1.7 (1.3-2.1) | 0.468 |
| **LVEF (%)** |  |  |  |
| Pre | 42 (13) | 50 (11) | 0.007 |
| Post | 46 (12) | 49 (9) | 0.209 |

**Figure 1.**
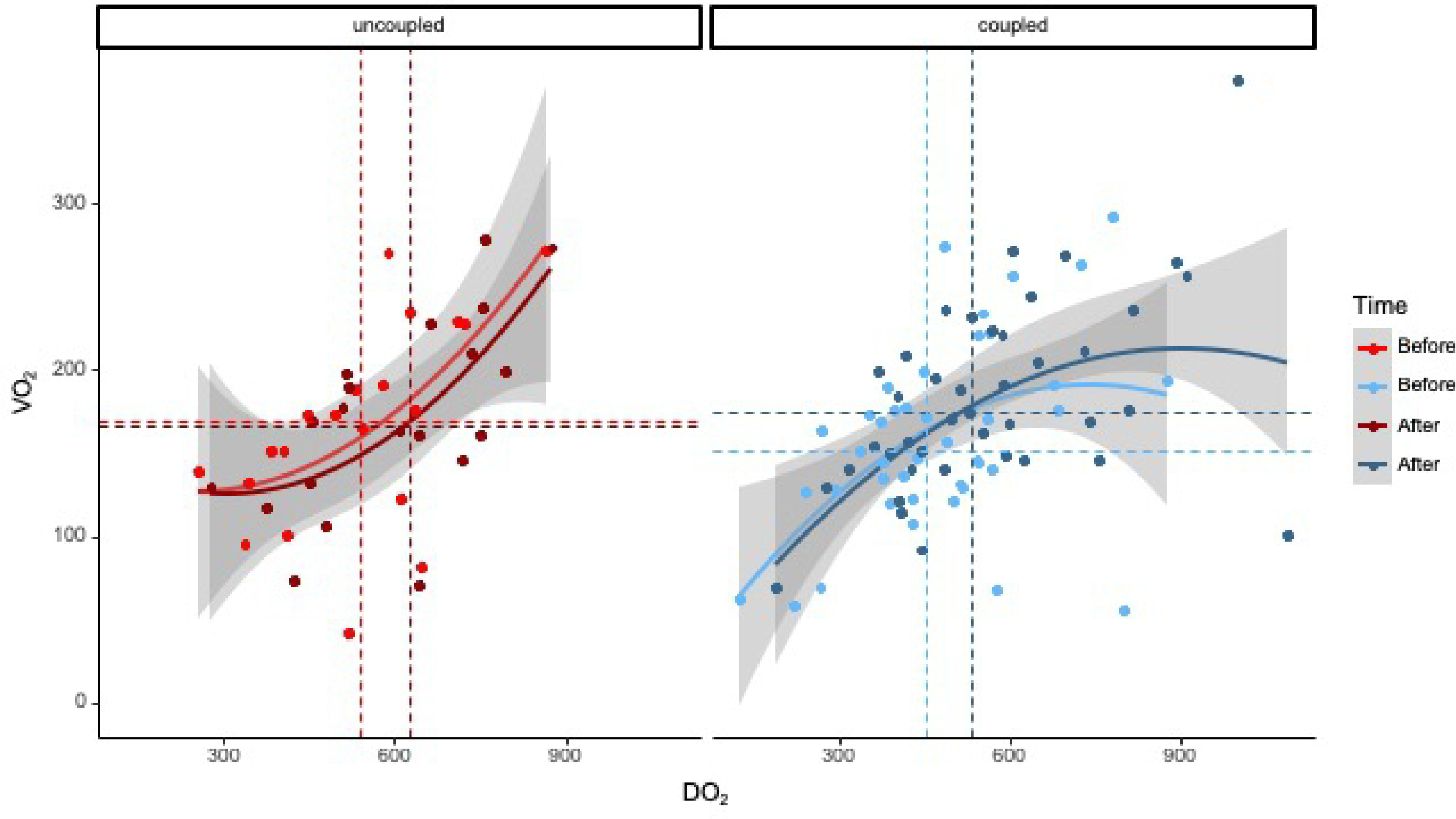
V-A coupling ratio according to VO_2_ change

### Effects of FC on systemic oxygenation parameters (Supplementary File 2)

At baseline, VO_2_ responders had lower VO_2_, gapCO_2_, and higher E_A_/E_V_ ratio, ScVO_2_ than VO_2_ non-responders. FC increased SAP, MAP, and SV in VO_2_ responders and non-responders. VO_2_ responders were characterized by an increase in SV and CO decreased TPRi, E_A_, and an increased SW/PVA ratio and gapCO_2_. E_V_ did not change. VO_2_ non-responders were characterized by an increase in SV, ScVO_2_, and a decreased HR.

### Effects of norepinephrine on systemic oxygenation parameters (Supplementary File 2)

At baseline, VO_2_ responders had lower SV, CO, SW/PVA ratio, VO_2_, and higher gapCO_2_, E_A_/E_V_ ratio than VO_2_ non-responders. Norepinephrine infusion increased SAP, MAP, CO and DO_2_ in both groups. VO_2_ responders were characterized by an increased SV and CO, SW/PVA ratio. VO_2_ non-responders were characterized by an increased E_A_, E_V_, ScVO_2_.

### Correlations between systemic oxygenation parameters (efficacy) versus E_A_, E_V_, E_A_/E_V_ ratio and SW/PVA ratio (efficiency) with the two therapeutic interventions

In the overall cohort, changes in VO_2_ were correlated with those in SW/PVA ratio (r=0.362, p=0.003), E_A_ (r=-0.446, p<0.001), E_A_/E_V_ (r=-0.256, p=0.046), CO (r=0.495, p<0.001), ScVO_2_ (r= −0.522, p<0.001), TPRi (r=-0.444, p<0.001). The baseline SW/PVA ratio was correlated with DO_2_ (r=0.339, p=0.004), VO_2_ (r=0.258, p=0.045), and gapCO_2_ (r=-0.304, p=0.017).

Baseline E_A_/E_V_ was predictive of VO_2_ responsiveness, with an area under the curve (AUC) [95% confidence interval (95%CI)] of 0.76 ([0.62-0.87]; p=0.001). With an AUC [95%CI] of 0.72 [0.59-0.85] (p=0.004), baseline ScVO_2_ was predictive of VO_2_ responsiveness. When analysing patients separately for FC or norepinephrine infusion, baseline E_A_/E_V_ was predictive of VO_2_ responsiveness in FC group (AUC: 0.77 [0.59-0.95] (p=0.008) and norepinephrine group (AUC: 0.74 [0.56-0.93] (p=0.045).

When using the principal component analysis, the 3 first principal components explained 61% of the variance (Supplementary file 2 and Figure 2). VO_2_ changes were significantly associated with the first (r=0.31) and the third component (r=0.52). E_A_, E_V_, E_A_/E_V_, and SW/PVA ratio were variables included in components associated to VO_2_ changes.

**Figure 2.**
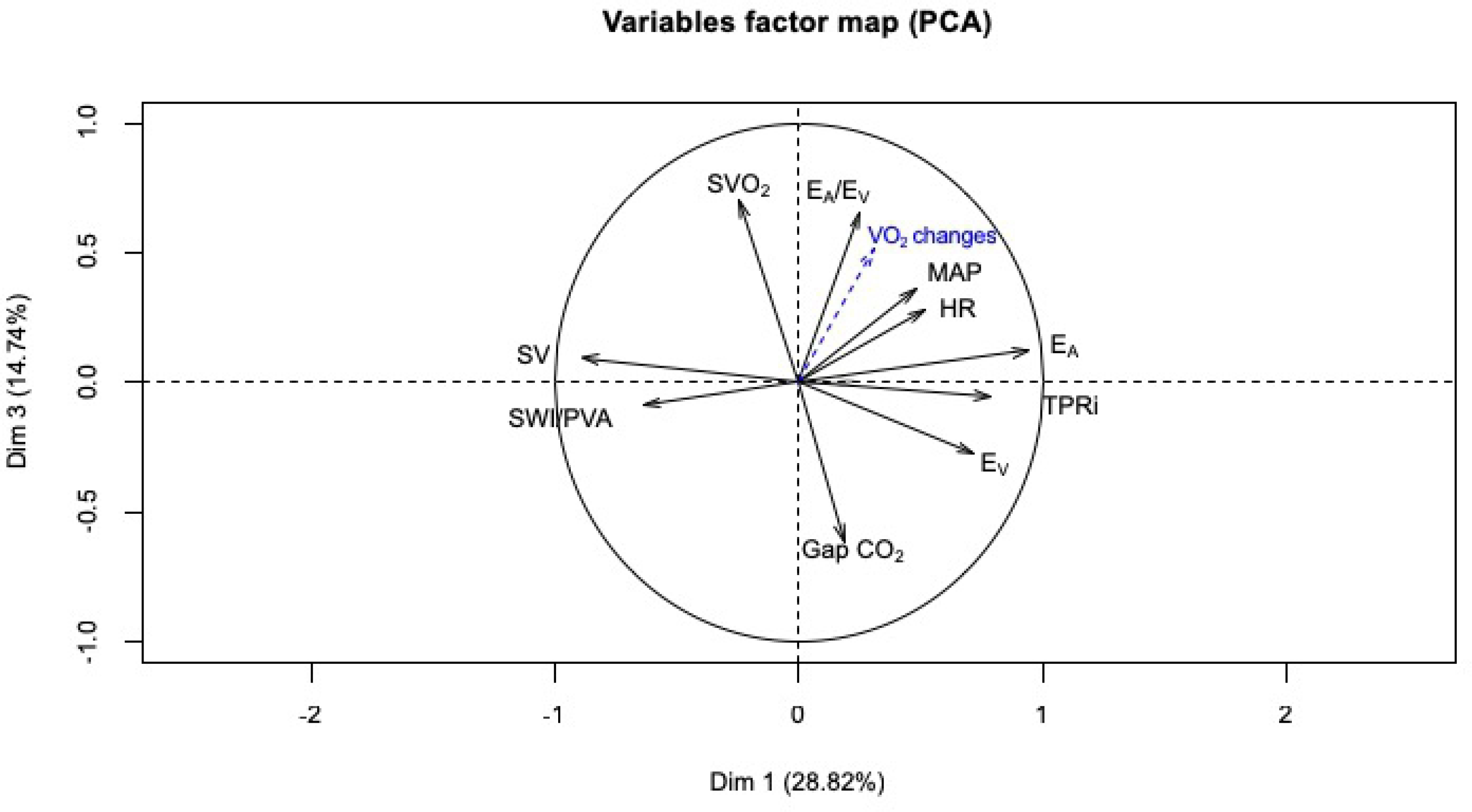
Variable factors map for the first and third component of the principal component analysis. The 10 more contributory variable are represented.

## DISCUSSION

The main results of the present study are as follow: (1) the majority of patients for whom FC or norepinephrine infusion increased VO_2_ had V-A uncoupling with lower SW/PVA ratios; (2) baseline E_A_/E_V_ and SW/PVA ratios were associated with perfusion parameters and VO_2_ changes independently of the therapeutic intervention used.

When analysing together FC or norepinephrine infusion, the only common profile is the increase in arterial pressure. VO_2_ responders have an increase in SV, CO and a decrease in TPRi. VO_2_ responder patients were uncoupled before interventions as they adapted to maintain tissue perfusion with a higher energetic cost for the same efficacy (preserving efficacy over efficiency); which was reflected by the lower E_A_/E_V_ ratio in VO_2_-responders. Equally, VO_2_-responder patients had significantly lower SW/PVA values before hemodynamic intervention, which were associated to perfusion parameters. We demonstrated the E_A_/E_V_ was part of components that explain VO_2_ responsiveness, and was independently associated with VO_2_ responsiveness. Both approaches credibly establish at least the statistical relevance of analysing V-A (un)coupling in patients with hemodynamic instability following cardiac surgery.

### A pathophysiological perspective on V-A coupling

Burkhoff and Sagawa have shown that mechanical efficiency is greatest when E_A_ = E_V_ (i.e E_A_/E_V_ ratio =1) (4, 6, 17). The patients of the present study were characterized by “normal” E_A_ but much lower than normal E_V_ values thus resulting in 78% patients having V-A uncoupling. Burkhoff and Sagawa have also shown that the mechanical efficiency of the heart is more sensitive to E_A_, especially when E_V_ is impaired, which is observed at baseline in the patients of the present study (4, 6, 17). “Sacrificing” efficiency to preserve efficacy for a limites period of time is a “physiological choice” observed in athletes (20). The patients with the highest V-A coupling ratio (uncoupled) had the lowest VO_2_ (20). The consequences of long term “sacrificing”, i.e. days for ICU patients are not known. For instance, the fact that catecholamine use is associated with increased mortality could be an example of long-term consequences of better cardiovascular performance with a high energetic cost (21).

Investigating the effects of two interventions on V-A coupling comes down to ask the question already raised many years ago: how effectively an increase in myocardial performance (an increase in SV) is transmitted to the peripheral circulation (22). This transmission may be mediated by the V-A coupling (22). In this respect, if the increase in cardiac performance is transmitted to the circulation, this should result into opening of new vascular beds, and if DO_2_ limits the VO_2_, this should result into an increase in VO_2._ This is exactly what our results demonstrate, linking the increase in cardiac performance with the peripheral circulation through the V-A coupling.

### Clinical relevance of V-A coupling in ICU patients

The clinical relevance of the statistical relationship between V-A (un)coupling and VO_2_ in the context of goal-directed therapy in critically-ill patients is still to be demonstrated. Septic shock is characterised by different profiles of V-A uncoupling (i.e different hemodynamic profiles) for which hemodynamic treatment may differ (23). The use of beta-blockers is an illustration of hemodynamic optimisation based on V-A coupling perspective (24, 25). Cardiologists have already integrated this hemodynamic approach in the treatment of chronic heart failure or arterial hypertension (9, 18, 25). Moreover, V-A coupling has been demonstrated as a factor limiting patients’ adaptability to effort (8, 20). Few studies have investigated the relationship between V-A coupling on one side and DO_2_ and VO_2_ on the other in ICU patients (10, 26). To the best of our knowledge, this is the first attempt that has specifically focused on VO_2_. Previous authors have studied the association of V-A coupling improvement and the time course of systemic oxygenation parameters in trauma patients (26, 27). Our results support their findings by demonstrating an association between V-A coupling, SW/PVA ratio, to perfusion parameters and further VO_2_ changes. One advantage of V-A coupling may be the fact that it can be non-invasively measured at bedside. On contrary to perfusion parameters, it does not need blood sample. The final clinical relevance of V-A (un)coupling for cardiac surgery patients will require well designed interventional trials such as the one publised by Borlaug et al that used LV afterload reduction (28).

### Potential limitations of the present study

The analysis of two therapeutics can make interpretation of the results difficult. The present objective was not to precisely analyze the individual effect of each therapy. Such demonstrations have been previously and extensenly studied (14). On the contrary, we would like to demonstrate that hemodynamic approach based on the V-A coupling makes it possible to dispense with the hemodynamic treatement and a detailed analysis of each parameters. The fact that the association between V-A coupling and perfusion parameters was demonstrated in the population as a whole and in each treatment group reinforces our results. As discussed, we believe that the effects of norepinephrine on VO_2_ may be due to its effects on CO and DO_2_ (29). The VO_2_/DO_2_ relationship is not linear. The VO_2_ responder group has lower value of VO_2_ that is below the value of the VO_2_ non-responder group, even after hemodynamic treatment. We believe the lower value of VO_2_ in responder group may have not introduce bias. These observations are in relation with the fact that the hemodynamic response was defined by VO_2_ changes. The methods used to calculate E_V_ and E_A_ can potentially be criticized because we did not use a high-fidelity ventricular pressure catheter (14). We calculated ESP from a radial artery signal, which may differ from the aortic pressure signal. However, radial artery pressure has been reported to provide a good estimate of ESP (30). Although it can be argued that estimation of ESP from the radial artery has not been fully validated, any error in this method would only affect the precision of absolute values of E_A_ and E_V,_ but not the E_A_/E_V_ ratio, as the error in end-systolic pressure would be similar. Despite these limitations, non-invasive evaluation of E_V_ and E_A_ was validated against the gold standard method, and has been used in cardiac surgery (5–7). In the present study, E_A_ and E_V_ must be considered to be approximations of E_A_ and E_V_. Despite these limitations, non-invasive evaluation was validated against the gold standard method, and have been used in cardiologic and cardiac surgical area (14, 18).

## CONCLUSIONS

In VO_2_ responders, V-A coupling was characterized by a high E_A_/E_V_ ratio (due to high E_A_ and low E_V_). Baseline E_A_/E_V_ and SW/PVA ratios were associated with VO_2_ changes independently of the hemodynamic intervention used. Measuring V-A coupling may offer a new perspective of hemodynamic optimisation in ICU by individualising hemodynamic treatment and by analysing both the efficacy and efficiency of hemodynamic interventions.

## Acknowledgements

All authors read and approved the manuscript.

The authors declare that they have no competing interests.

No external funding was received.

The study received no specific financial support. This work was supported by the Department of Anaesthesiology of CHU de Amiens, Amiens, France.

